# SPECTRE-Plex: an automated, fast, high-resolution-enabled approach for multiplexed cyclic imaging and tissue spatial analysis

**DOI:** 10.1101/2025.01.30.635753

**Authors:** Michael D. Anderson, Abigail Plone, Jeffrey La, Madison Wong, Krishnan Raghunathan, Jocelyn A. Silvester, Jay R. Thiagarajah

## Abstract

Mapping the spatial organization of tissues is critical to understanding organ biology in health and disease. Developments in multiplexed antibody-based, fluorescence labelling methods have provided unique insights into tissue microenvironments. However, many current methods have a variety of limitations that reduce their practical utilization. To address this, we developed SPECTRE-Plex; an end-to-end solution based on a series of methods that significantly improve speed, automation, and resolution of cyclic multiplex immunofluorescence imaging.

---

Describing the spatial context of how proteins and cells organize, interact and change is critical to understanding tissue and organ biology in health and disease. Several different high dimensionality spatial biology approaches for tissues have emerged that detect and measure either mRNA transcripts (e.g. MERFISH, Slide-seq)^1,2^ or protein (e.g CODEX, cyCIF)^3–7^.

Detection and localization of proteins within tissues allows assessment of cellular function, local communication, and accurate mapping of tissue state. Imaging-based protein methods center around cognate antibody binding to targets of interest, with either fluorescence or mass-spectroscopy-based detection. To detect multiple targets in the same tissue, most multiplexed protein techniques are based on cyclical, iterative antibody binding^8^. Discrimination of individual targets between cycles can be achieved in a variety of ways such as fluorescence dye inactivation (t-cyCIF, IBEX), barcoding (CODEX) or antibody elution^3,5,9,10^. Each of the major methods has relative advantages and disadvantages varying by application, ease of use, and resource investment. For example, while CODEX enables highly amplified and specific signal identification, the need for specific oligonucleotide conjugation substantially increases cost (for commercially available targets/panels) and/or time (for custom antibody conjugation and validation). Antibody elution / stripping methods have advantages in linear signal amplification but can be hampered by incomplete antibody removal and loss of tissue morphology after a finite number of cycles^8^. Dye inactivation methods based on directly conjugated fluorescent antibodies are attractive because of the large array of cheap, validated commercially available antibodies, and relatively simple procedures and reagents. However, current implementations of dye inactivation methods have limitations related to cost, speed, resolution, and fluidics.

We therefore developed an end-to-end system, based on a series of novel or optimized methods. **S**patial **P**hoto-inactivation **E**nhanced **C**yclic **T**arget **RE**solved multi**Plexi**ng (SPECTRE-Plex) broadly encompasses three modules. Firstly, an optimized upright water dipping objective based imaging and fluidic system, including custom solutions for repeatable, iterative image capture. Secondly, a novel microfluidic imaging chamber system incorporating a refractive index matching optical window that enables imaging of tissues at a variety of resolutions including the use of high NA (>1.0) objectives in conjunction with repeated solution changes. Thirdly, a new fast, non-gas producing dye inactivation method that reduces cycle times. We show application of this system for a 22-plex, 7-round panel in healthy and diseased human duodenal tissue, including downstream analysis. In addition, we highlight additional applications of the method including direct tissue-based measurements of antibody binding kinetics for quantitative validation and assessment of staining conditions.

Figure 1a and Supplementary 1 outline the overall system design including custom optics with a water dipping objective, small volume fluidics system and python-based imaging and device control. The system is designed so that slide-mounted samples undergo staining, washing, imaging and dye inactivation in situ allowing efficient and rapid repetition of cycles (Figure 1b). Overall, the method significantly reduces experimental time. As an example, a 7-cycle run can be completed in 7 hours as compared to 14-36 hours for most manual or semi-automated cyclical methods (Figure 1b and Supplementary 3). Tiled imaging in place is enabled by a highly repeatable motorized stage. Integrated control of imaging, positioning and fluidics is achieved by custom python and μ-manager software (Supplementary 8-10).

**Figure 1:**
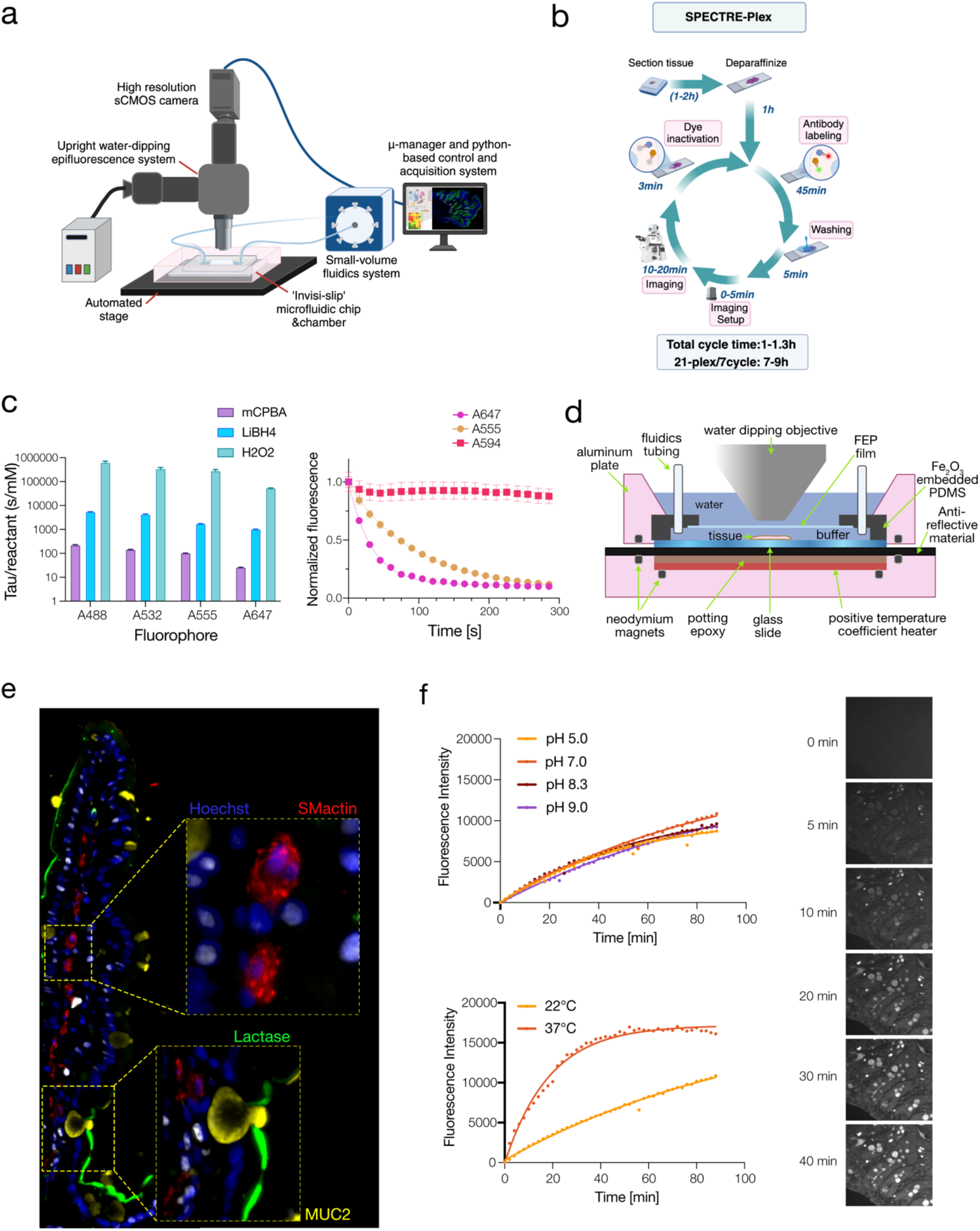
**(a)** Schematic of SPECTRE-plex system. **(b)** Overview of the cyclical steps and experimental times for SPECTRE-plex tissue staining. **(c)** Kinetics of dye inactivation of Alexa 488, 532, 555, and 647 showing time constant (*τ*, s/mM) for m-CPBA, LiBH_4_, and H_2_O_2_ (left) and time course of m-CPBA dye-inactivation for susceptible fluorophores Alexa 555 and 647 and non-susceptible Alexa 594 (right). **(d)** Schematic for imaging chamber system including microfluidic (‘invisi-slip’) device. **(e)** High magnification image of human duodenum using 60x, 1.2NA objective. **(f)** Antibody binding association of anti-MUC2 IgG on duodenal tissue with alteration of solution pH (top), and temperature (bottom) with representative images of antibody fluorescence (right).

To further reduce cycle times, we examined the chemistry underlying inactivation of cyanine and rhodamine-based fluorophores (Supplementary 4). This led to the discovery of meta-chloroperoxybenzoic acid (m-CPBA) as a potentially more efficient catalyst than previously used dye-inactivation agents. m-CBPA is orders of magnitude faster against a set of commonly used conjugated fluorophores, than lithium borohydride (LiBH_4_) or hydrogen peroxide (H_2_O_2_) (Figure 1c, Supplementary 5). In addition, m-CPBA does not generate oxygen in solution overcoming a major issue with LiBH_4_ and H_2_O_2_ for imaging and the application of microfluidics. We found that m-CPBA does not cause appreciable deterioration of tissue quality (tested up to 10 cycles) and the lack of bubble formation allows easy implementation of small volume fluidic flow control.

Imaging resolution and magnification of current automated cyclical fluorophore staining methods is limited by the use of air objectives (usually NA≤0.8) and/or glass chamber windows which introduce refractive index mismatching. To solve this issue, we developed a PDMS-based microfluidic device (‘invisi-slip’) with an embedded fluorinated ethylene propylene (FEP) window enabling aberration free imaging as the FEP polymer has the same refractive index as water and can be coupled to water-based objectives. This device is incorporated into an integrated imaging chamber system which includes ports and tubing for fluids and temperature control (Figure 1d). For whole tissue imaging and to reduce the number of imaging tiles we use a large field of view (FOV) 16x 0.8 NA water dipping objective. To illustrate the wide range of objectives that can be used we also carried out SPECTRE-plex runs using a 60x 1.2NA water dipping objective which allowed imaging of target proteins at sub-cellular resolution (Figure 1e).

The in-situ nature of the SPECTRE-plex design allows for simultaneous imaging and conjugated antibody labelling on tissues. This enables experiments assessing the kinetics of antibody-target association directly on tissues. To illustrate this, we tested the binding kinetics of a specific antibody (anti-Muc2 IgG) and the effect of altering solution pH or temperature. As shown in Figure 1f, at 22°C, anti-Muc2 exhibited a one phase association with a mean half-time of 50mins, which was not altered by solution pH but was significantly reduced (t_½_=13mins) by increasing solution temperature to 37°C. This capability of the SPECTRE-plex system is relevant to a variety of applications including tissue specific validation of antibodies and rapid assessment of therapeutic antibody or small molecule binding.

To demonstrate the use of our system, a 22-plex image data set was generated using duodenal tissues from children with active celiac disease and age-matched healthy controls. Figure 2a shows representative registered and stitched whole tissue SPECTRE-plex images including markers of specific cell types such as epithelial cells (Na^+^K^+^-ATPase), T-cells (CD3d), enteroendocrine cells (ChromograninA), and myofibroblasts (smooth muscle actin) among others (Supplementary 7). A custom image analysis pipeline involving adaptive segmentation and cell coordinate mapping (Figure 2b, Supplementary 12) enables several downstream analyses. Cell compositional analysis (Figure 2b,c,d) revealed differences between healthy and celiac disease tissue including increases in intestinal enteroendocrine cells, immune cell expansion and increased spatial cellular heterogeneity. Celiac disease tissues showed relative expansion of proliferating (PCNA+) epithelial cells with focal high-density areas and increases in cellular density within non-epithelial compartments (Figure 2e,f,g). Analysis of tuft cells, a lineage of epithelial cells involved in luminal nutrient and pathogen sensing^11^ showed differences in both distribution and clustering between healthy and celiac disease tissues (Figure 2h,i). Neighborhood cell-cell analysis for pEGFR+ tuft cells indicated increases spatial association with both proliferating and differentiated epithelial cells (Figure 2j).

**Figure 2:**
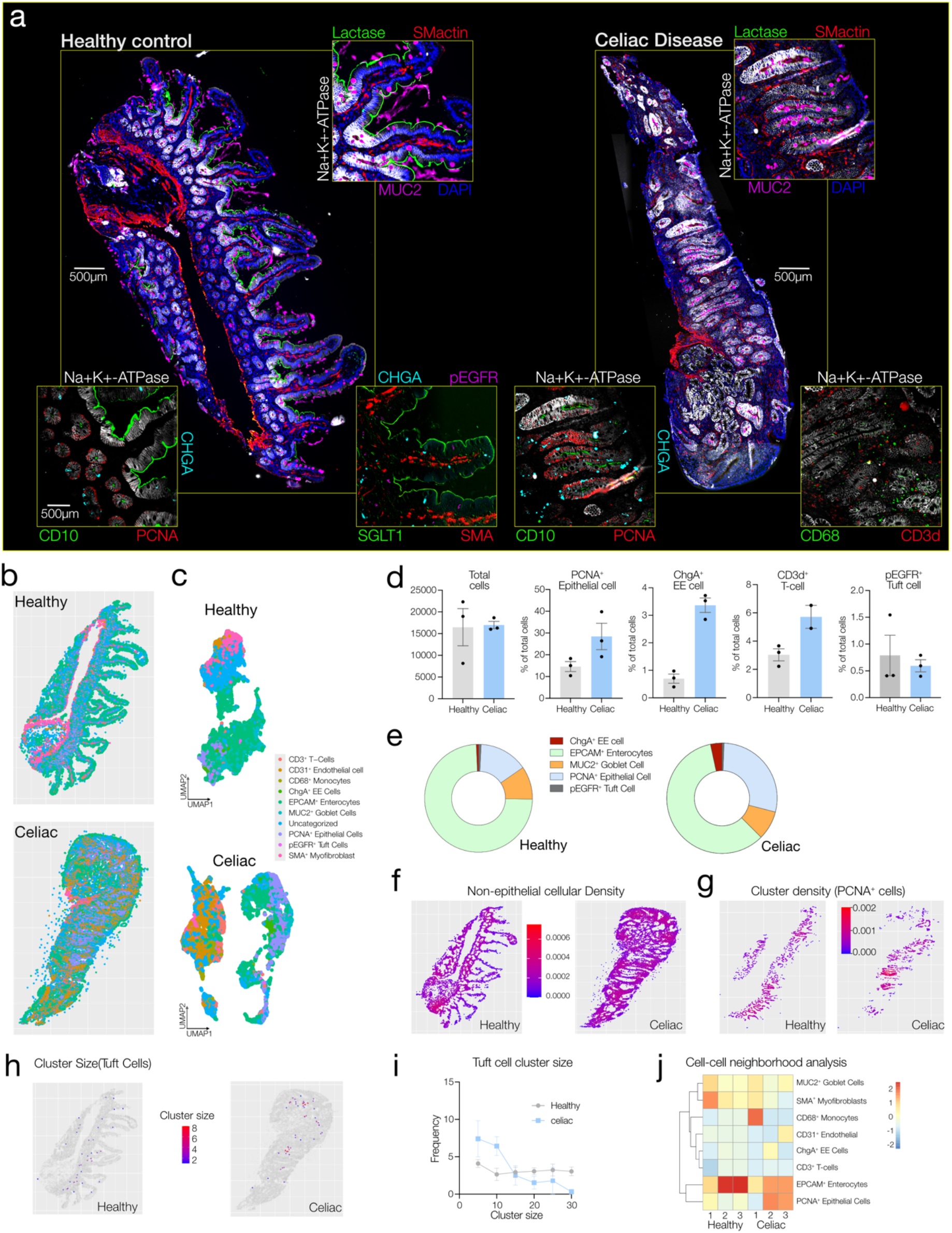
**(a)** Example tiled, stitched, processed whole section tissue images from healthy control duodenum and active celiac disease. Main image showing lactase, mucin-2 (MUC2), smooth muscle actin (SMactin), sodium-potassium ATPase (Na^+^K^+^-ATPase) and nuclei (DAPI), with inset target antigens as indicated. **(b)** Cell-type spatial map of healthy and celiac disease duodenum. **(c)** Uniform Manifold Approximation and Projection (UMAP) for spatial clustering of cell-types. **(d)** Mean cell numbers and cell percentages for healthy versus celiac disease tissue (n=3). **(e)** Mean epithelial sub-type cell percentages for healthy versus celiac disease tissue. **(f)** Example cellular density analysis for non-epithelial cells. **(g)** Example spatial cluster density analysis for PCNA+ cycling epithelial cells. **(h)** Example spatial cluster size analysis for pEGFR+ tuft cells. **(i)** Mean cluster size frequency for pEGFR+ tuft cells (n=3) **(j)** Cell-cell neighborhood association analysis for pEGFR+ tuft cells.

Future improvements include the addition of a multi-slide configuration to enable high throughput experiments and screening, and addition of confocal optics to enable three-dimensional sample imaging. Although SPECTRE-Plex was primarily designed for dye-inactivation multiplex protein profiling, the imaging and microfluidic modules can be readily applied to other tissue-based imaging applications including RNA profiling, barcoding, and other use cases such as quantitative tissue binding studies.

## Supplementary Information

### Supplementary 1: Imaging chamber design

**Figure S1:**
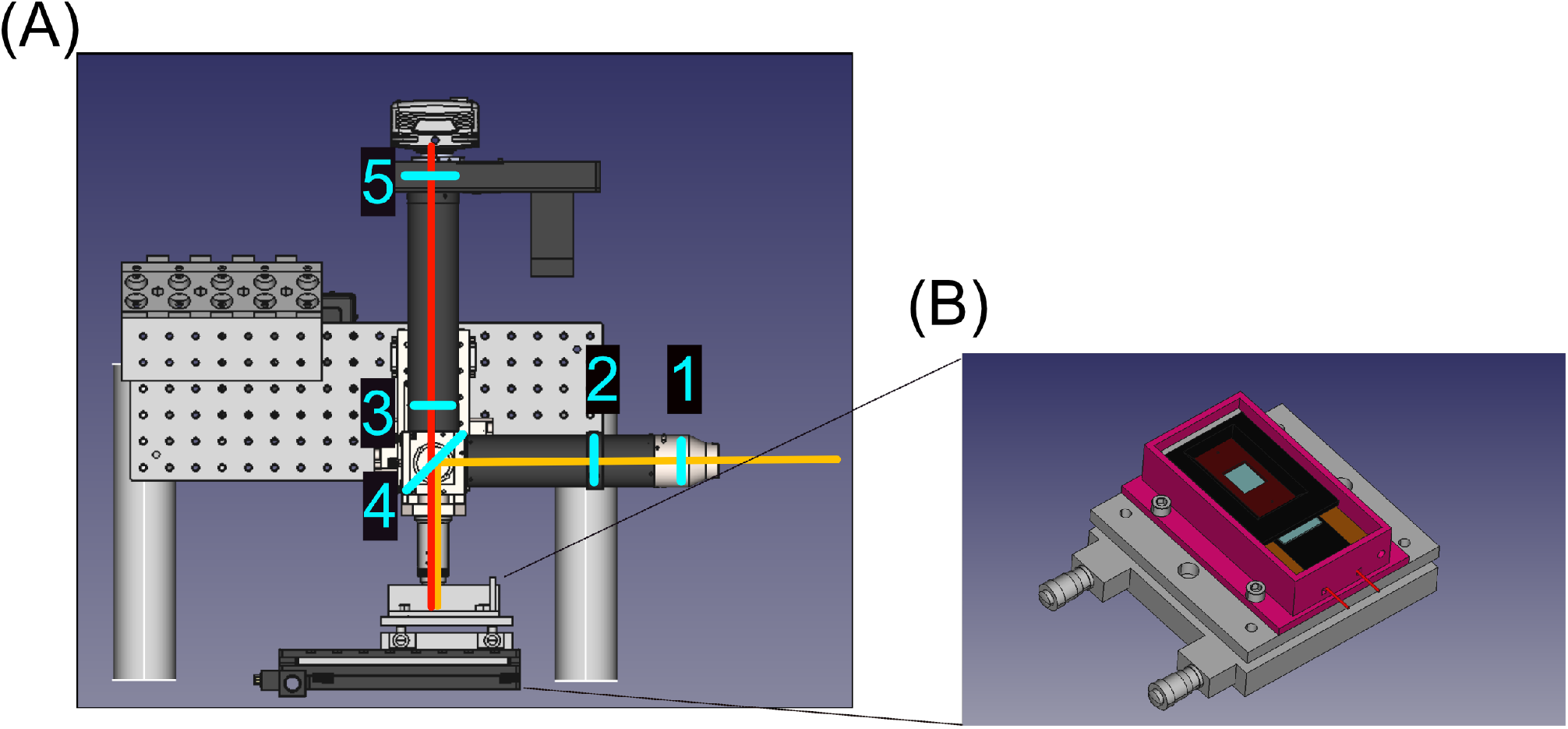
**(A)** 3D rendering of system with light path and lens superimposed. The elements in the light path are an (1) 38mm focal length collimating, liquid light guide coupling lens, (2) excitation tube lens, (3) pentapass dichroic, (4) Nikon 16x 0.8NA water dipping objective, (5) emission tube lens, (6) emission filters in an emission filter wheel and a photometric IRIS 15 camera. **(B)** 3D rendering of imaging chamber with slide into alignment jig, invisi-slip fluidic device on slide and pressure plate in place over invisi-slip device.

The optical train for the SPECTRE-Plex is a custom design using the modular infinity microscope platform from Applied Scientific Imaging (ASI). This is an upright 4F widefield configuration with a Kohler illuminated, liquid light guide fed excitation path and an additional extra-long Nikon tube (265mm vs 200mm) to increase magnification to eliminate vignetting. The excitation source is a Lumencor light engine (Spectra III) containing integrated excitation filters, with a Photometrics IRIS 15 camera and Nikon CF175 series 16x 0.8NA water dipping objective. The fluidic system comprises of an Elveflow OB1 mk4 air pressure regulator with inline flow meter, Elveflow multiplexing valve (12×1) and a custom 3D printed device to hold and organize the individual Eppendorf reservoir adapters. The stage is a combination of an ASI XY OE 1250 stage and a TechSpec tilt stage. The XY stage has a high travel range and is highly repeatable for positioning. The tilt stage allows for fine tuning of the chamber tilt to be parallel with the camera imaging plane.

The main body of imaging chamber is 3D printed out of PLA and has two sets of glued in neodymium magnets. One set of magnets is to interface with the pressure plate, while the other set is to aid in the alignment of the Invisi-Slip microfluidic device. A custom positive temperature coefficient (PTC) heater from Thermo-heaters LLC is adhered to main chamber body and a hard-set potting epoxy is poured over the heater and the magnets and pressed flat until it set. This encases the magnets and heater in epoxy, isolates them from the water in the chamber and enables good thermal conductivity. This also provides a very flat surface for the slide to rest on and seals the pores in PLA of the chamber body to make it watertight. The next layer, a polymer film called Maxi Black (Acktar) abrogates reflection in the system by absorbing a wide band of frequencies and preventing spectral scattering. A PLA 3D printed alignment jig allows reproducible placement of both the slide’s positions in the chamber and placement of the pressure plate over the slide. Finally, the pressure plate was milled out of aluminum (Protolabs) and anodized to prevent any oxidation and contains sockets for neodymium magnets. The magnets were encased in a poured UV curable resin. Specific details of parts and construction are available on request.

### Supplementary 2: Protocol and Methods

#### Standard SPECTRE-Plex Protocol

##### Reagents

- Histoclear (ThermoFisher Scientific, cat no. 50-329-48)
- Ethanol (Decon Laboratories, cat no. 2716)
- PBS (Corning, cat no. 21-040-CV)
- Hoechst (ThermoFisher Scientific, cat no. H3570)
- LiBH4 (Strem 9300397)
- H2O2 (Sigma 216763)
- m-CPBA powder (Sigma 273031)
- bovine serum albumin (Sigma-Aldrich, cat no. A6003-25G)
- Triton-X-100 (Sigma-Aldrich, cat no. X100-100ML)
- Citrate stock solution (Abcam, cat no. ab93678)
- DAKO Diluent (Agilent Dako, cat no. S080983-2)
- Water

#### Buffer Preparation

<u>Bleach base solution:</u> 24mM NaOH was prepared by diluting 10M NaOH stock in PBS.

<u>Blocking Buffer:</u> 0.1% Triton-X-100 and 5% bovine serum was added to PBS.

<u>Antigen Retrieval Buffer:</u> The Citrate stock solution was diluted 100-fold in PBS.

<u>m-CPBA Bleach Solution</u>: m-CPBA stock was prepared by dissolving m-CPBA powder in 100% ethanol at 1M concentration or 172mg/mL. This stock was used for up to 30 days. For the working bleach solution, the stock was diluted 1:100 in bleach base solution.

<u>LiBH_4_ Bleach Solution:</u> Dissolve the appropriate amount of LiBH4 in distilled water for desired concentration (10mM for bleach kinetics and 50mM for bubble generation demonstration). The solutions were made from powder opened within the previous 30 days and used immediately after preparation.

<u>H_2_O_2_ Bleach Solution:</u> 30% H2O2 (Sigma 216763) was added to bleach base solution to make 4.5% H2O2.

<u>Antibody Dilutions:</u> Prepare 500 uL antibody dilutions from DAKO Diluent containing 1:25k dilution of Hoechst and desired antibodies for a given staining round.

<u>Phosphate-Citrate Buffer for pH experiments:</u> 100mM concentration was used for all pH experiments. Buffers were made by adding appropriate amount of Sodium Phosphate Dibasic Dihydrate and citric acid. Sodium Hydroxide and Hydrochloric acid were used for fine pH adjustment. Ionic strength was calculated to be within 5% for each chosen pH value.

##### Slide Preparation

1. For deparaffinization, place slides in a 60°C bead bath for 15 min.
2. Incubate with 100% Histoclear for 5 min at RT. Repeat this step twice with fresh Histoclear each time for 5 min and then 10 min at RT.
3. For rehydration, incubate slides in decreasing concentrations of Ethanol.
  a. First, incubate in 100% Ethanol for 5 min at RT. Repeat this step with fresh Ethanol for 5 min at RT.
  b. Incubate in 95% Ethanol for 5 min at RT. Repeat this step with fresh Ethanol for 5 min at RT.
  c. Incubate in 75% Ethanol for 5 min at RT.
  d. Incubate in 50% Ethanol for 5 min at RT.
  e. Incubate in tap water for 5 min at RT.
4. Move slides into a plastic Coplin jar and pour Antigen Retrieval Buffer in until tissue on slides is completely covered.
5. Submerge the Coplin jar into a boiling water bath for 20 min.
  - The set-up is a 1L beaker of boiling water on the heating plate with a stripette on the top to which the holder is attached with a binder clip.
6. Remove the Coplin jar from water bath and let cool at RT for 20-40 mins.
7. Wash with 1x PBS for 5 min. Repeat this step with fresh 1x PBS.
8. Pour Blocking Buffer into the Coplin jar and let sit at RT for 1 hour.
9. Store in 1x PBS at 4°C until ready to stain.

##### Multiplex Set up

10. When ready to start SPECTRE-plex run, stain slide with 1:5k Hoechst dilution for 5 minutes. Give slides a quick rinse with 1x PBS.
11. Prepare the antibody dilutions (described above).
12. Mount a slide into the base of the chamber and place the fluidic device on top. Carefully lower the top plate making sure the fluidic device stays in desired placement. Insert the tubing onto either side of the fluidic device.
13. Check if a seal has been created between the slide and fluidic device by running PBS through tubing.
  a. Note: If PBS does not flow, then the seal has not been created. Remove the tubing and reseat the top plate if this occurs and test again.
14. Move the chamber so it is positioned underneath the objective. Open the Micro-manager software and lower the objective.
15. Locate the sample using the DAPI channel. Check if the sample is level.
16. Use the MicroMagellen plugin to define a surface.
17. Give the project folder a name and start the program. All custom python programs are available in our github page.

#### Comparison of different bleach solutions

The following fluorophores were evaluated for their propensity to photobleach under different bleach solution:

1. Alexa-488 (Invitrogen prod# A28175, 1mg/mL)
2. Alexa-532 (Life Technology A11002, 2mg/mL)
3. Alexa-546 (Phallodin)
4. Alexa-555 (Invitrogen A21428, 2mg/mL)
5. Alexa-568 (Life Technology A11031, 2mg/mL)
6. Alexa-594 (Chromogranin)
7. Alexa-647 (Life Technologies A21246, 2mg/mL)

Bleach solutions were prepared as described. All experiments were conducted as triplicates in a 96 well format using in a Tecan Spark multi-well plate imaging system in inverted fluorescence mode with the monochromator positions chosen to have 25nm bandwidths centered on excited and emission peaks for each fluorophore. Each fluorophore was diluted at 1:1000 in PBS and with the appropriate concentration of each of the different bleach solution with the bleach solution without any fluorophores as their respective controls. The average fluorescence intensity normalized to the control was measured every 15 seconds for 10 minutes. The resulting kinetic data was fit in Graphpad Prism using a double exponential decay model.

#### Antibody Staining Kinetics

The kinetics experiment was conducted on the SPECTRE-Plex system and modified from the standard SPECTRE-Plex imaging protocol. A slide stained with DAPI was loaded onto the microscope’s stage and washed with PBS. For these experiments, a fixed exposure time and illumination intensity (50% power and 50ms exposure time for both Alexa-488 and Alexa-647) was used. An initial image was taken as the background autofluorescence image and then the fluidic system was used to dispense the stain solution. Next, every 2 minutes, the system would execute the stardist^1^ recursive autofocus algorithm (as described in the autofocus section) and each channel imaged. A final image was taken 90 minutes after a PBS wash. Temperature and pH were modulated for these kinetic experiments. For the pH experiments, Phosphate-citrate buffer was used due to its large buffering range. Temperature was modulated using an embedded heater in the imaging chamber. For these experiments, the heater was then turned on for 1.5 hours in order to hit a stabile temperature which was measured by an infrared thermometer prior to antibody staining.

Images were processed by subtracting the background, zero-time frame and the average pixel intensity of the tissue region was calculated for each frame. A binary tissue mask was generated by thresholding the final background-subtracted image via Otsu’s method. An exception was made for the pH 5.0 for MUC2 data as the signal was too low for Otsu’s method to threshold accurately. In this case, the mask was set by manually tracing out 3 small regions of goblet cells in the image. The temporal change in average pixel intensity data was fit to a single exponential function in Graphpad Prism.

#### Assessment of fluidic bubble generation

H_2_O_2_, LiBH_4_ and m-CPBA were mixed at normal working concentrations (4.5%, 50mM, 10mM respectively) in 15mL tubes, vortexed and left to sit for 20 minutes for observing the propensity of each solution to form bubbles.

### Supplementary 3: Comparison of SPECTRE-Plex with manual or semi-automated tCYCIF

**Figure S3:**
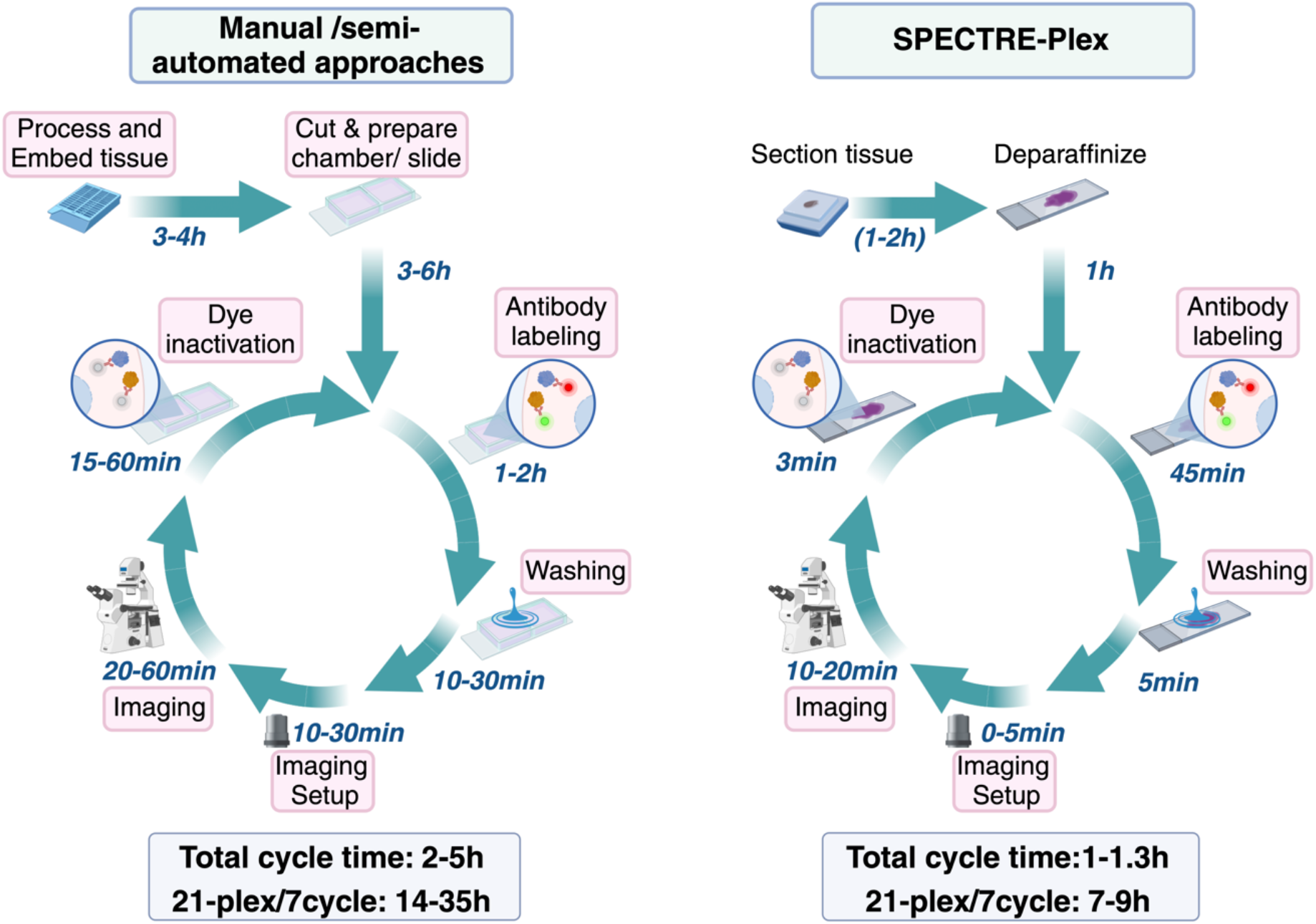
Estimated total experimental time for a 7 cycle multiplex run based on manual and semi-automated tCYCIF protocols (left)^2^ versus the SPECTRE-Plex system (right).

### Supplementary 4: Chemical reaction for dye inactivation

**Figure S4:**
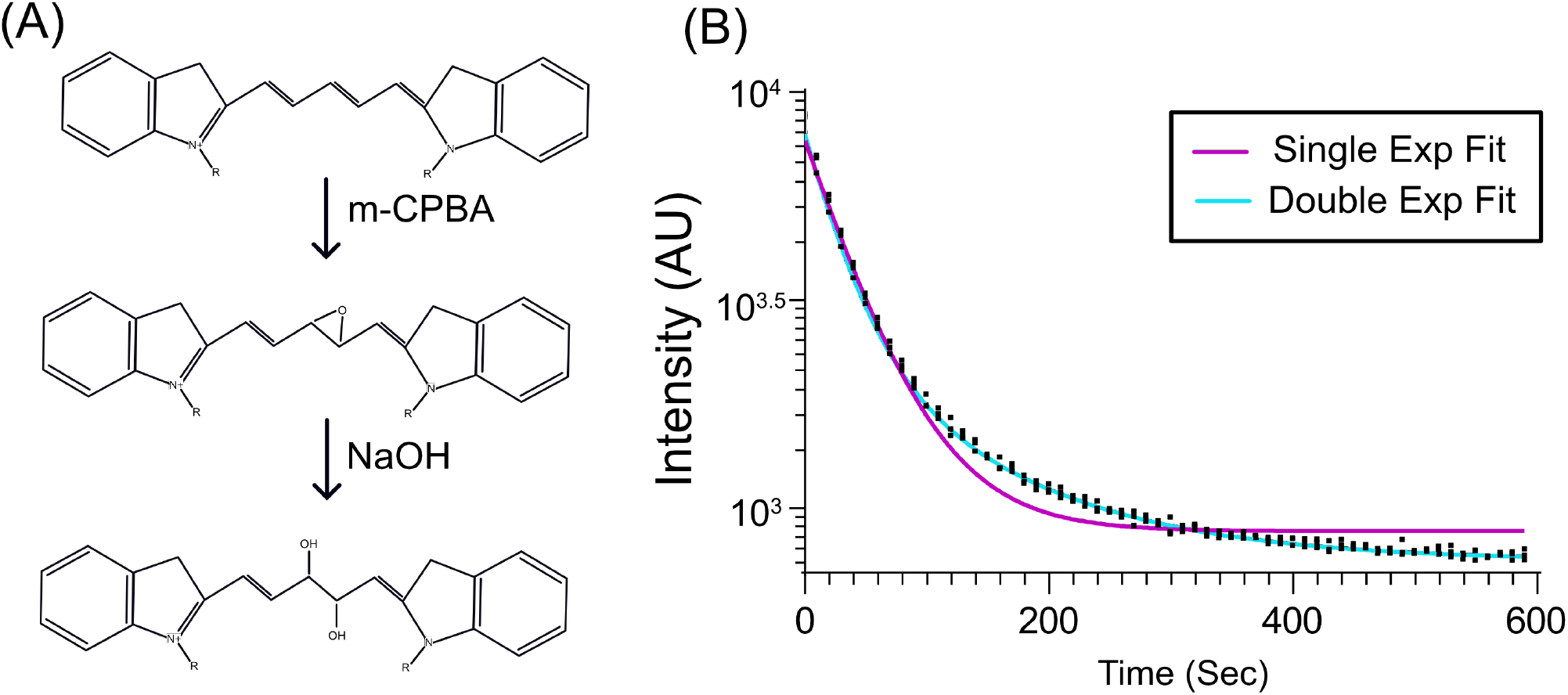
**(A)** The proposed chemical reaction is epoxidation of the alkene groups via the Prizelhaev reaction followed by the formation of vicinal diols via the addition of NaOH. **(B)** Bleaching of Alexa-488 bleaching in the presence of 10mM meta-chloroperoxybenzoic acid (m-CPBA) and 20mM NaOH were analyzed to kinetic model based on either single or double exponential fitting. m-CPBA is a more efficient catalyst of the Prizelhaev reaction in comparison to hydrogen peroxide.

The kinetic data supports a sequential 2-part reaction. We observe that cyanine-based and rhodamine-based fluorophores are susceptible to bleaching.

### Supplementary 5: Comparison of mCPBA over LiBH4 and H2O2 for dye inactivation and oxygen production

**Figure S5:**
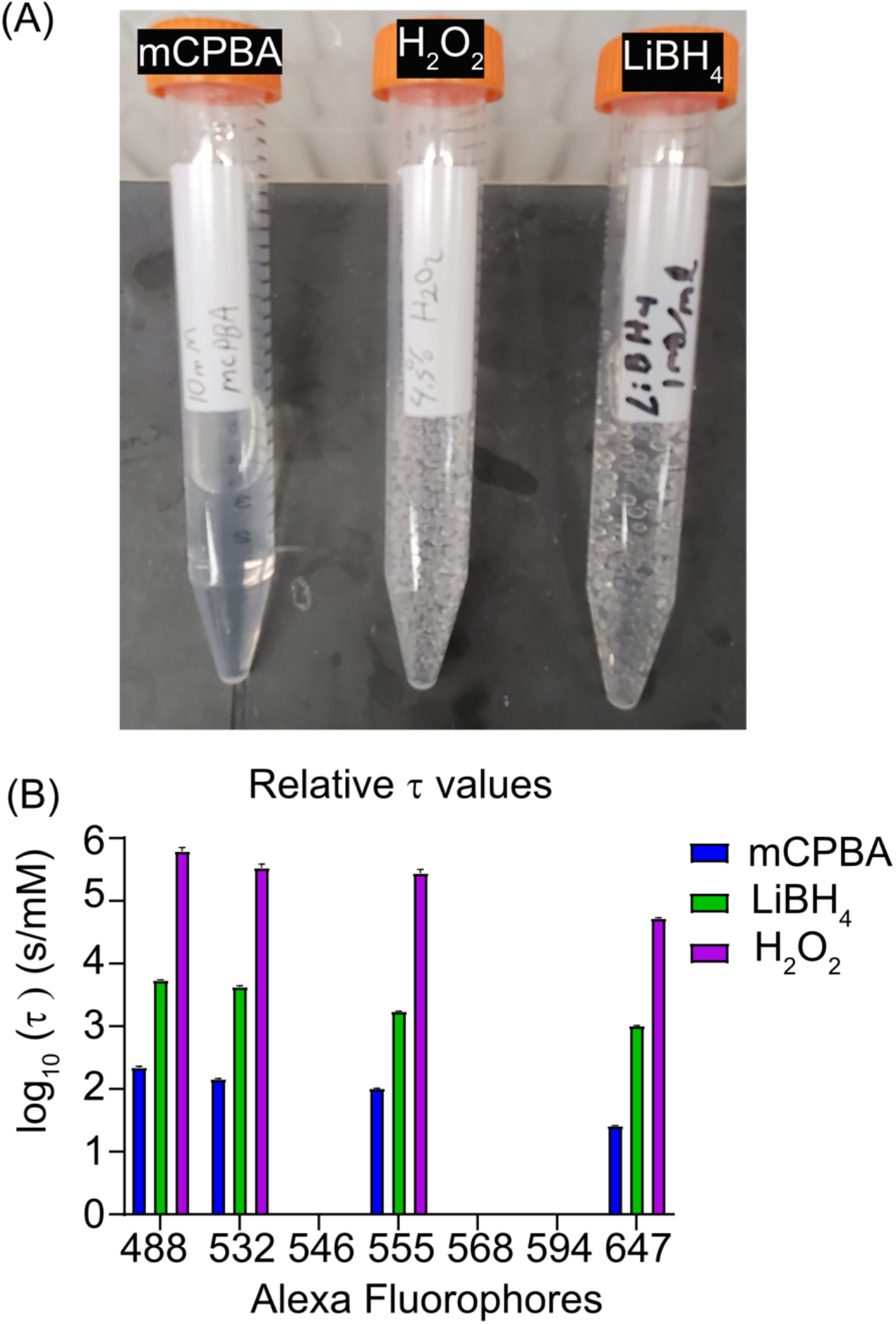
**(A)** Falcon tubes containing 10mM mCPBA, 4.5% H_2_O_2_ and 1mg/ml LiBH_4_ showing gas production over 20 minutes. **(B)** Comparison of *τ* values for bleaching

mCPBA has three advantages over LiBH_4_ and H_2_O_2_ for dye-inactivation and multiplex immunofluorescence applications. Firstly, mCPBA unlike both LiBH_4_ and H_2_O_2_ does not produce air bubbles for up to 24 hours on incubation with target fluorophores (Fig S. Absence of air bubbles is critical for small volume microfluidic applications as these can rapidly accumulate in any closed system either in tubing or in the imaging chamber. This leads to sub-optimal and variable image quality as well as variability in fluid flow control for solution exchange. Secondly, mCPBA has much higher chemical and solution stability than LiBH_4_ or H_2_O_2_ which are known to be relatively labile. In our experience, working concentrations of mCPBA continue to have sufficient efficacy for up to 36 hours and a 1M stock solution of mCPBA in ethanol is stable for 30 days. Thirdly, dye inactivation by mCPBA is orders of magnitude faster compared to LiBH_4_ and H_2_O_2_. For all the antibodies that we have tested, mCPBA was at least 10 times faster in bleaching than LiBH_4_ and H_2_O_2_ at the recommended working concentrations.

### Supplementary 6: Invisi-slip microfluidic device

**Figure S6:**
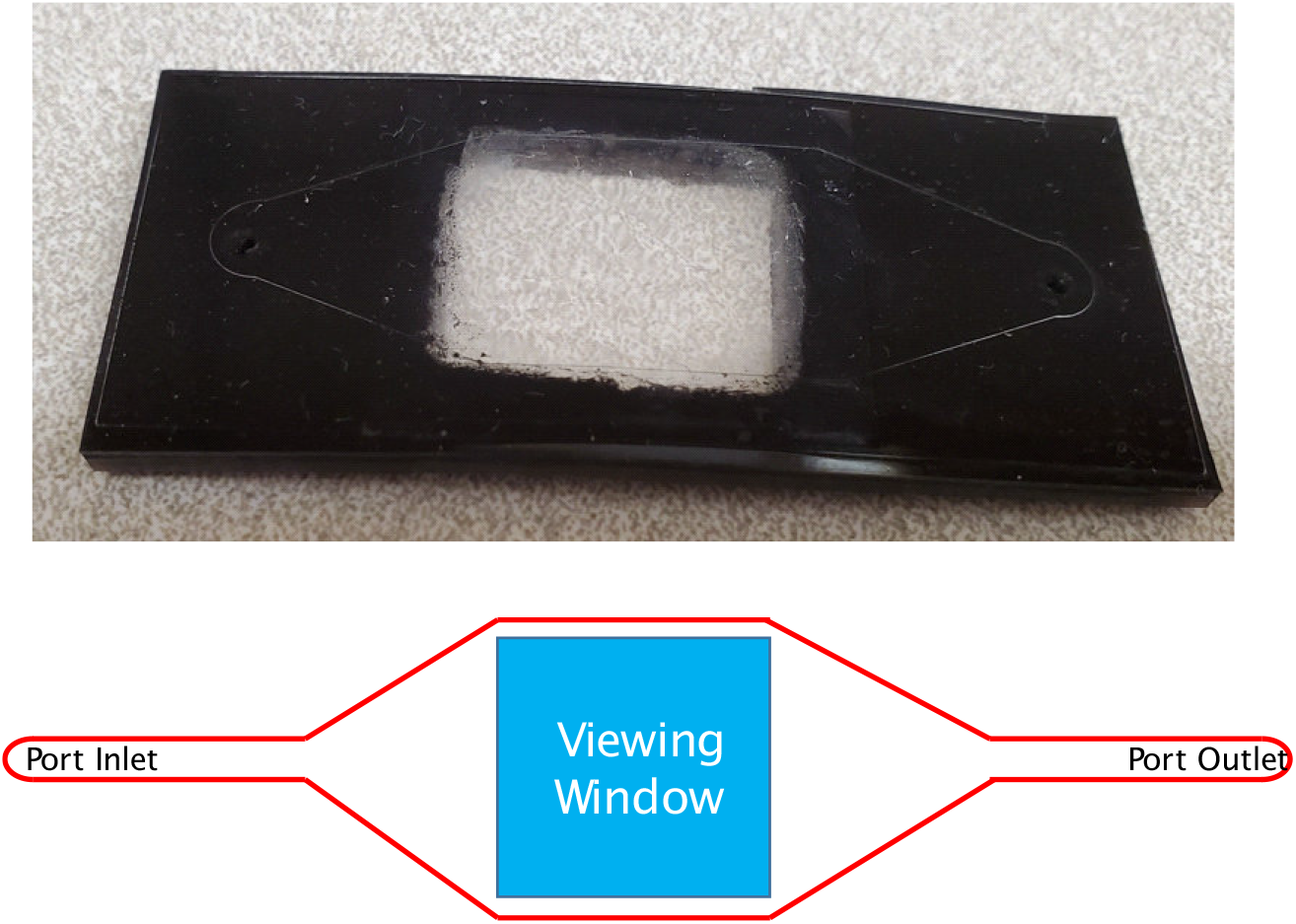
Overall design of Invisi-slip microfluidic device

Cover-slipping is an important engineering hurdle to overcome for automating multiplexing microscopy assays as the act of removing a coverslip, while trivial by hand, would require a robotic arm system to accomplish in an automated system. Most commercial automated platforms integrate these coverslips into the device that seals around the sample of choice, but this is often not readily adaptable for use in other systems. In addition, the use of coverslips and the consequent refractive index mismatching that occurs between fluid, glass and air results in inherent limitations to the numerical aperture (NA) and magnification of the imaging objective and therefore limitations to resolution. The Invisi-slip system circumvents these issues by using a Fluorinated ethylene propylene (FEP) polymer viewing window embedded in a PDMS device as a backbone. The PDMS backbone can be manufactured in any microfluidic device manufacturing facility that uses silicon wafer molds. The device is not plasma treated and is completely reusable. A magnetic pressure plate is used to seal the device to a slide. The 6 neodymium magnets in the pressure plate and 6 in the chamber below them with a 2mm gap between them, easily gives enough sealing force to have fluid move 1000uL/min within the device. The FEP viewing window can be made to be very large to accommodate large tissue sections or tissue microarrays. In our case, the window has dimensions of 18mm x 21mm. The internal volume of the device is approximately 120uL which allows the system to use reagent quantities that align with microliter volume fluidics systems. Overall, this system is a reusable, flexible solution that allows for aberration free imaging as the FEP polymer has the same refractive index as water and therefore can be coupled to water immersion or dipping objectives. Furthermore, the small distance from the FEP film window to tissue sample, allows for the use of high NA (>1.0), or high magnification objectives.

### Supplementary 7: List of antibodies used

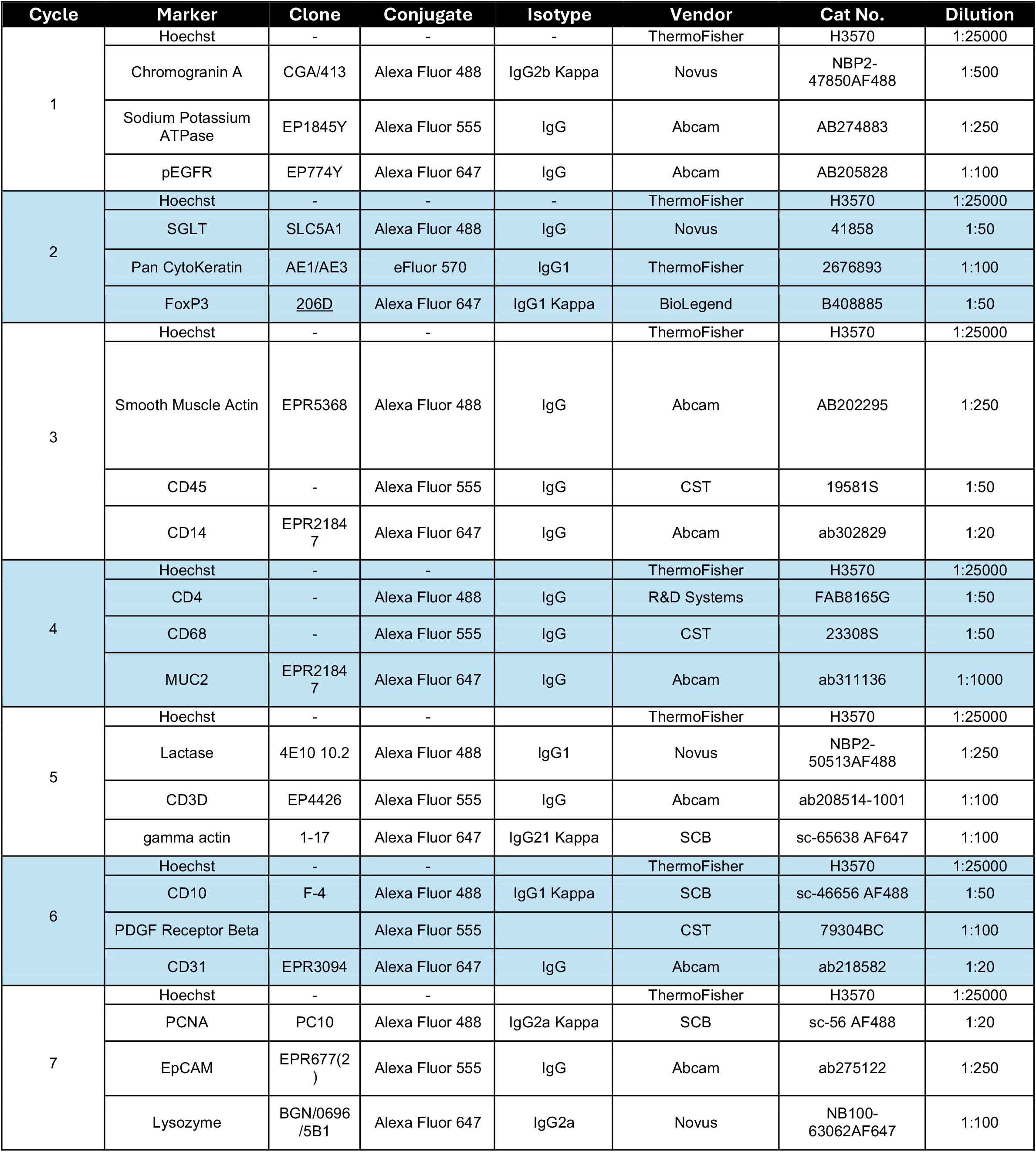

### Supplementary 8: Acquisition Outline

**Figure S8:**
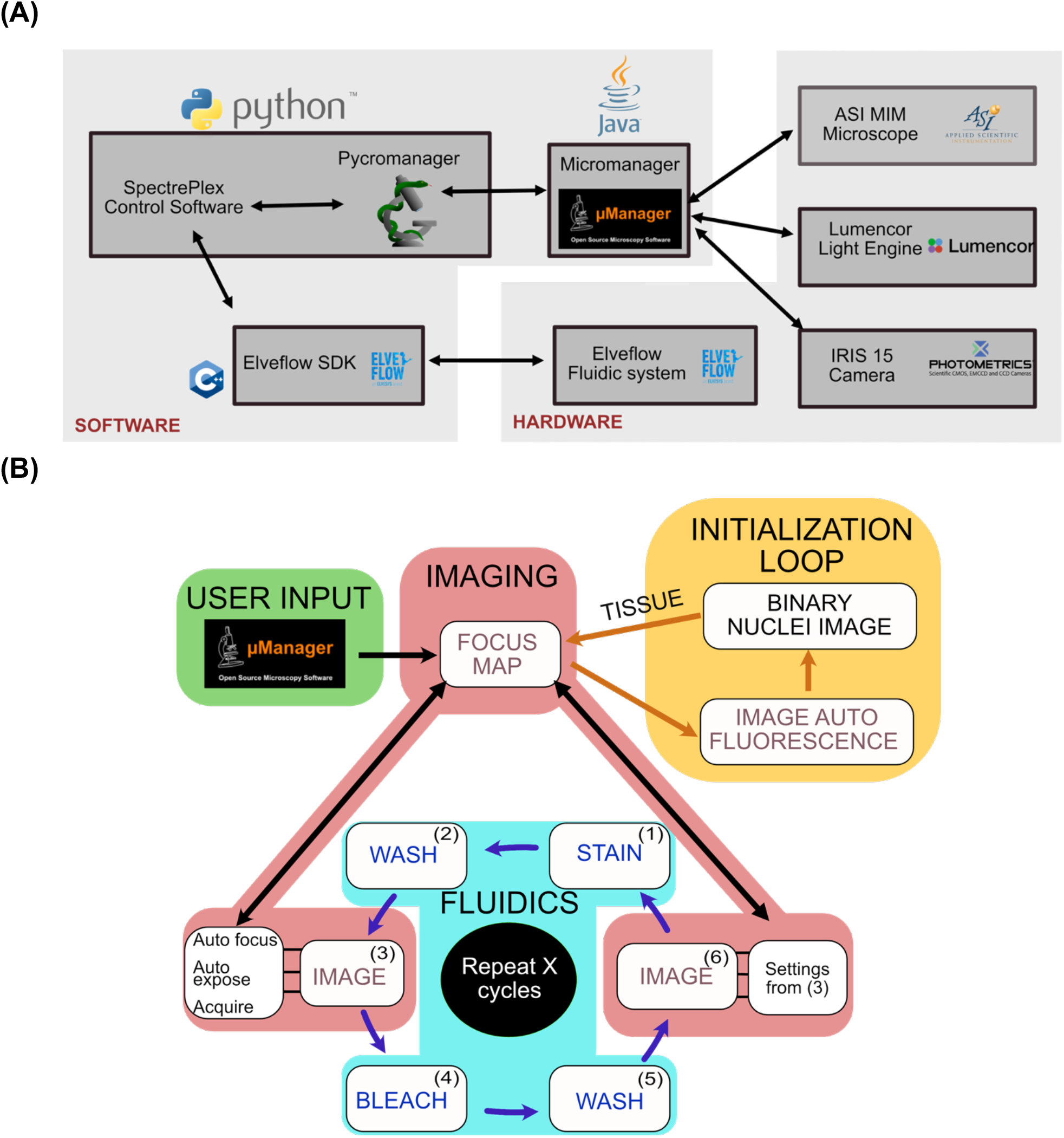
**(A)** An overview of the hardware and software components and their interconnectedness is outlined. **(B)** A visual guide to the overall acquisition process

An initial acquisition is performed to capture autofluorescence with default exposure times. The DAPI images are segmented as a binary image using Stardist^1^ and run through a size exclusion filter. The focus map is updated with this information for each tile to image only the tiles where tissue exists. Next, in the cycle process, stain is pumped into the chamber and incubated for 45 minutes, washed with PBS for 2.5 minutes and the slide was imaged. The next step in the cycle is the imaging, which has three parts (a) an auto focus step (b) an auto exposure step which updates the focus map and (c) an acquisition step to acquire the images. Following this is the dye inactivation step in which the imaging chamber is incubated with the dye inactivation solution for 3 minutes, followed by 2 minutes of washing. The fluorophore inactivated images were acquired using the exact same settings as the stained images. All code is available in Github at the following link: https://github.com/mdanderson03/SpectrePlex.git

### Supplementary 9: Autofocus

Brenner scores are used as a metric for determining the relative focus of images^3^. While this works well for ideal samples, we found that it had two major limitations. Firstly, in field of views (FOVs) where there is minimal tissue, the standard Brenner scores is highly weighted towards any bright region. Secondly, the standard Brenner score takes the difference between pixels that are two pixels away from each other in the x axis, i.e. *I*_*x*+2,*y*_ − *I*_*x*,*y*_. To practically improve the results for tissue-based applications, the Brenner score metric can be optimized by changing the number of pixels skipped. For a nuclei (DAPI) stain channel, we found that a pixel skip size of 17 to be optimal to maximize the Brenner score metric. To resolve the issue of minimal tissue in the FOV, we calculated a Brenner score constrained to only the tissue-positive region using a binary mask defined by the DAPI stain and using a standard StarDist network^1^. Given that Brenner score is weighed by bright spots, we have observed that the presence of fluorophore aggregates often leads to erroneous focus maps. We circumvented this by using the log of the image when calculating our modified Brenner score.

For an image with n x n pixels with *I*_*x*,*y*_ being its intensity at coordinates (x,y) and a binary image mask where *B*_*x*,*y*_ = 1 *or* 0 if nuclei are present at the coordinates (x,y), the focus scores will be calculated as the following:

Standard Brenner Score: 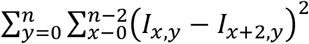

Modified Brenner Score: 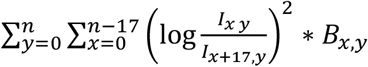

First, a 7-slice z stack with 2 µm step sizes in the DAPI channel is acquired. When the acquisition phase of the next cycle starts, the auto focus program uses the previous cycle’s DAPI images to determine on a per tile basis the slice that has the highest modified Brenner score. It then calculates the Δ*Z* between the current focus maxima with previous cycle’s maxima to center the stack around the current maxima and appends that value to the focus map. We observe that the average drift going from cycle to the next is on the order of 2-4 µm.

**Figure S9:**
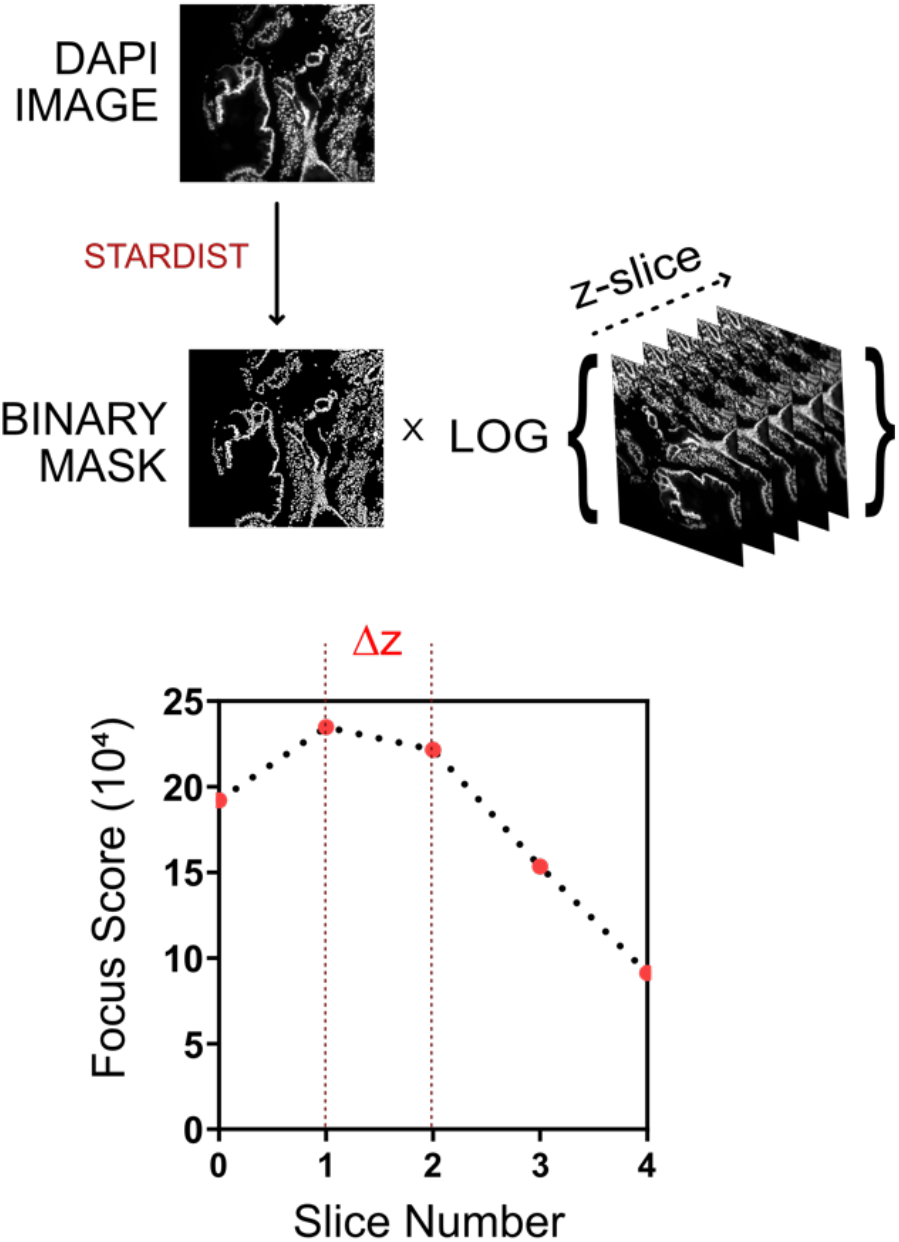
Overview of SPECTRE-plex autofocus routine using a optimized Brenner Score.

### Supplementary 10: Autoexposure

**Figure S10:**
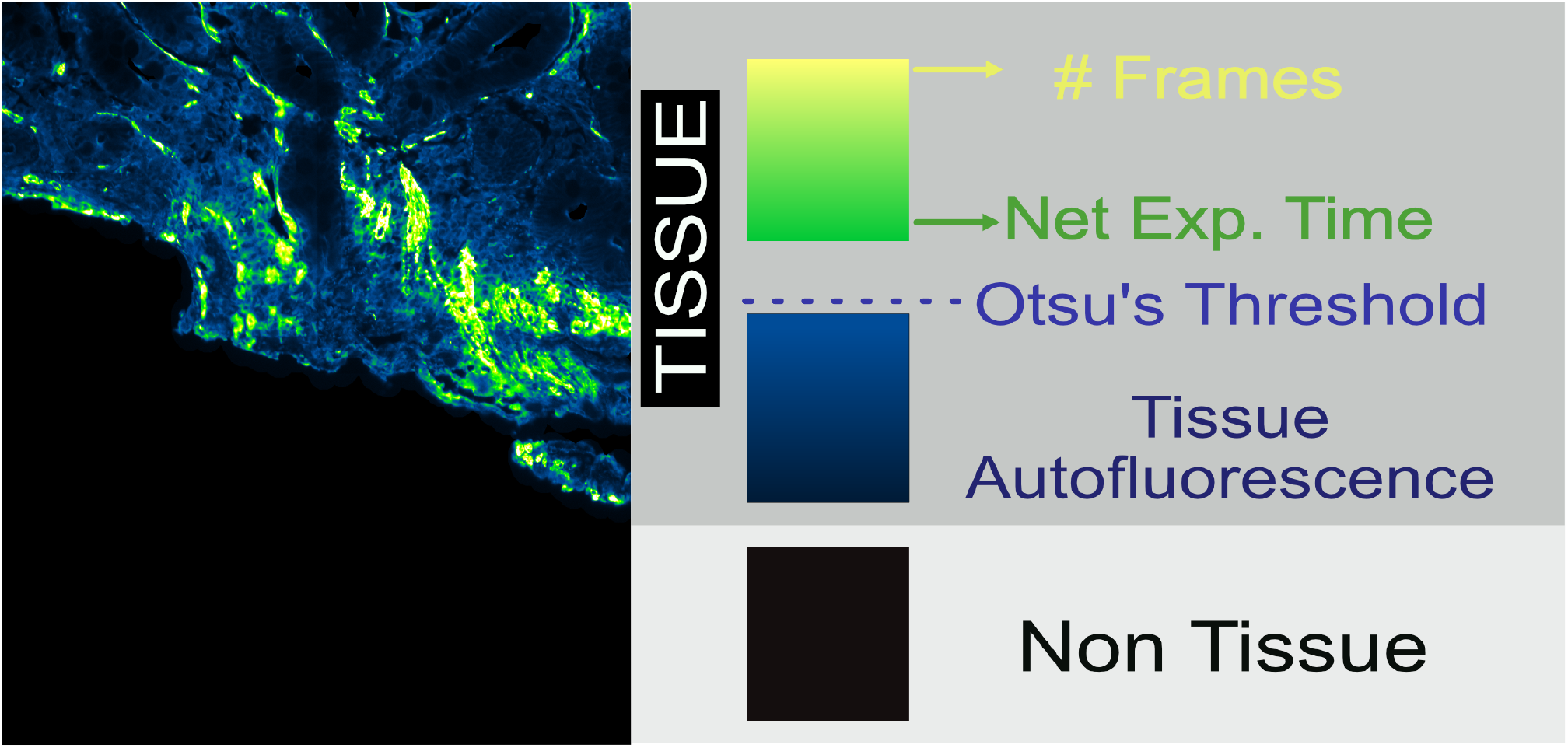
Overview of autoexposure routine.

The objective of auto exposure is to achieve an optimal signal to noise (SNR) ratio for each fluorophore stain. Here, the percent dynamic range occupied (PDRO) which measures the percentage of the dynamic range in the intensity histogram normalized to the total dynamic range was used as an approximation for SNR. Autoexposure was determined as follows.

First, we use a binary mask to specifically identify the tissue regions. Signal intensity in these tissue-associated pixels can contain autofluorescence or actual stain which can be distinguished using Otsu’s thresholding. From the subset of pixels which contain actual stain, the top and bottom 1% of the pixels by intensity are removed. These pixels are then used to determine the exposure time and the number of frames used for signal averaging.

1. 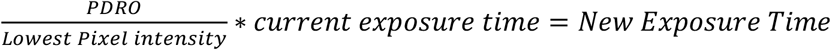
2. 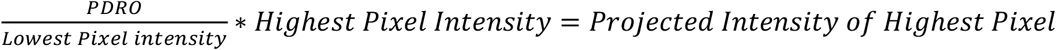
3. 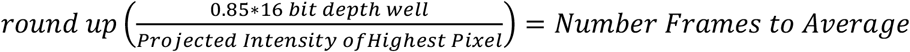
4. 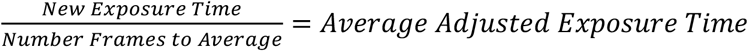

This method ensures that the dimmest pixels are adequately exposed, and the brightest pixels are not saturated. We have observed that while this method generally works well for most tissues, the presence of fluorophore aggregates tends to result in very small, calculated exposure times and underexposure of the actual stain. Similarly, very weak fluorophore staining tends to result in excessively high exposure times. To minimize the downstream effects on total imaging time that these scenarios can induce during automated operation of the SPECTRE-Plex system, we have implemented hardcoded strict lower and upper limits for the exposure time for any given fluorophore.

### Supplementary 11: Post-processing Pipeline

**Figure S11:**
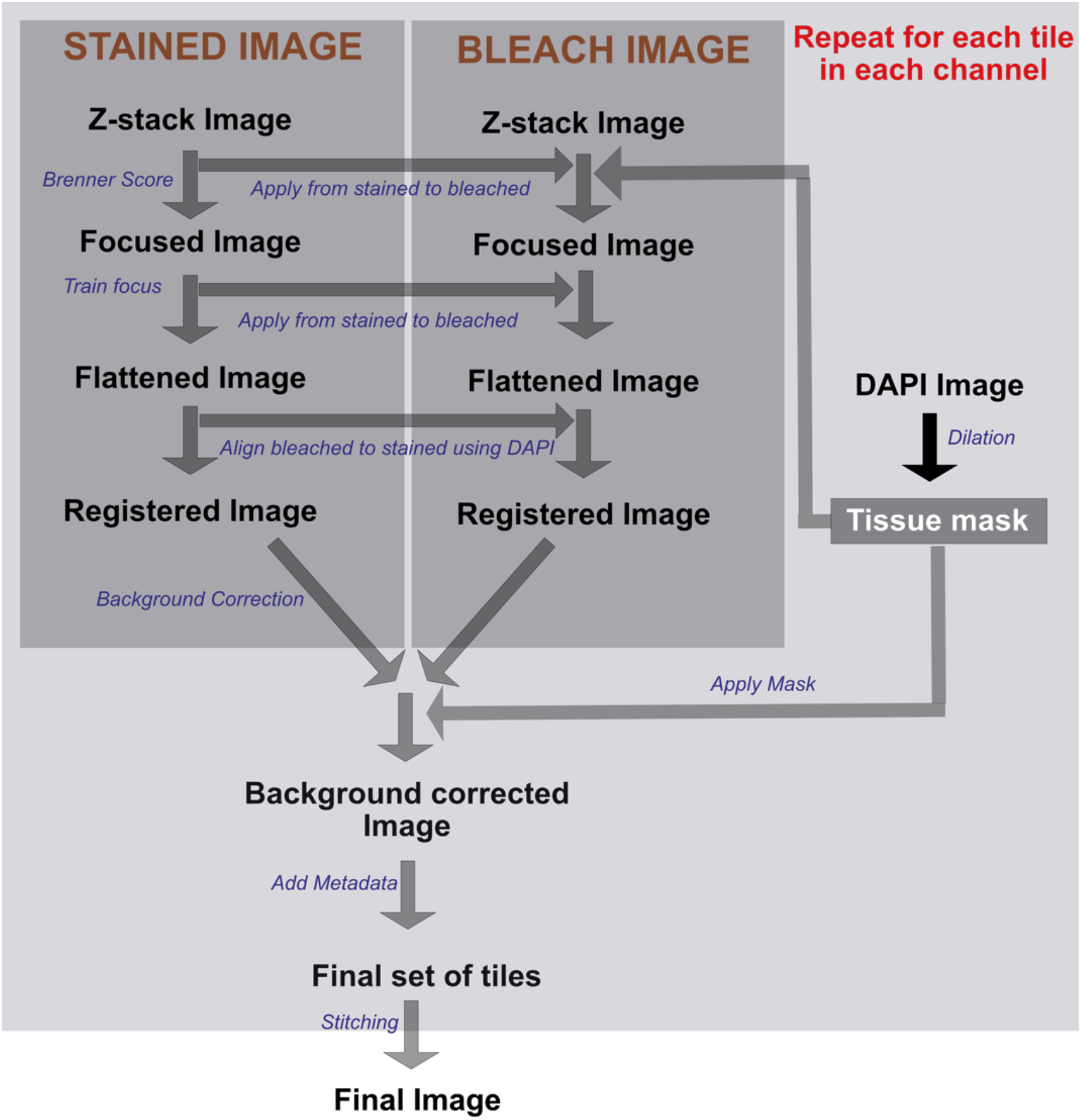
Overview of image processing pipeline

Image processing of SPECTRE-plex datasets occurs in 3 major categories: i.) determination of the infocus image, ii.) illumination normalization, and iii.) removal of background as outlined in Figure S11. Our standardized acquisition involves obtaining a z stack for each image area tile. This ensures that if the stain has a slightly different in focus location in z or if the DAPI offset is inaccurate for that specific tile, in focus information will still be captured. The modified Brenner score (see Supplementary 9) with a skip parameter of 10 instead of 17 is used to assess which z slice in the stack is in focus. The equivalent image in the dye-inactivated (‘bleach’) stack is obtained by using the same slice index as used for the stained z-stack. Next, the stained images that contain tissue regions are used to train a BaSiC^4^ flat field correction surface. This is applied to each stained and bleached tile. Since the bleached and stained images are gathered as part of different acquisition events, they may be slightly misaligned. Pystackreg^5^ is a python port of a turbo stack reg plugin from ImageJ and is used to register the bleached image to the stained image and output a bleached image following addition of a displacement vector. In practice this is generally only 1-2 pixels so no cropping is needed. Since the bleached image was acquired with the exact same setting as the stained, we subtracted the bleached image from the stained image. While this tends to leave some residual background, the majority is removed and additional processing such as a rolling ball background subtraction can be used if desired. After all tiles are processed, they are then packaged into tiff stacks and populated with metadata in such a way that they can be directly inputted into McMicro^6^. The Ashlar module within the McMicro pipeline which registers cycles and stitches tiles together is used to generate a registered and stitched output file. Lastly, a binary tissue mask is generated from the DAPI channel and multiplied by the registered image set to remove any non-tissue signal to output a finalized dataset.

### Supplementary 12: Analysis

**Figure S12:**
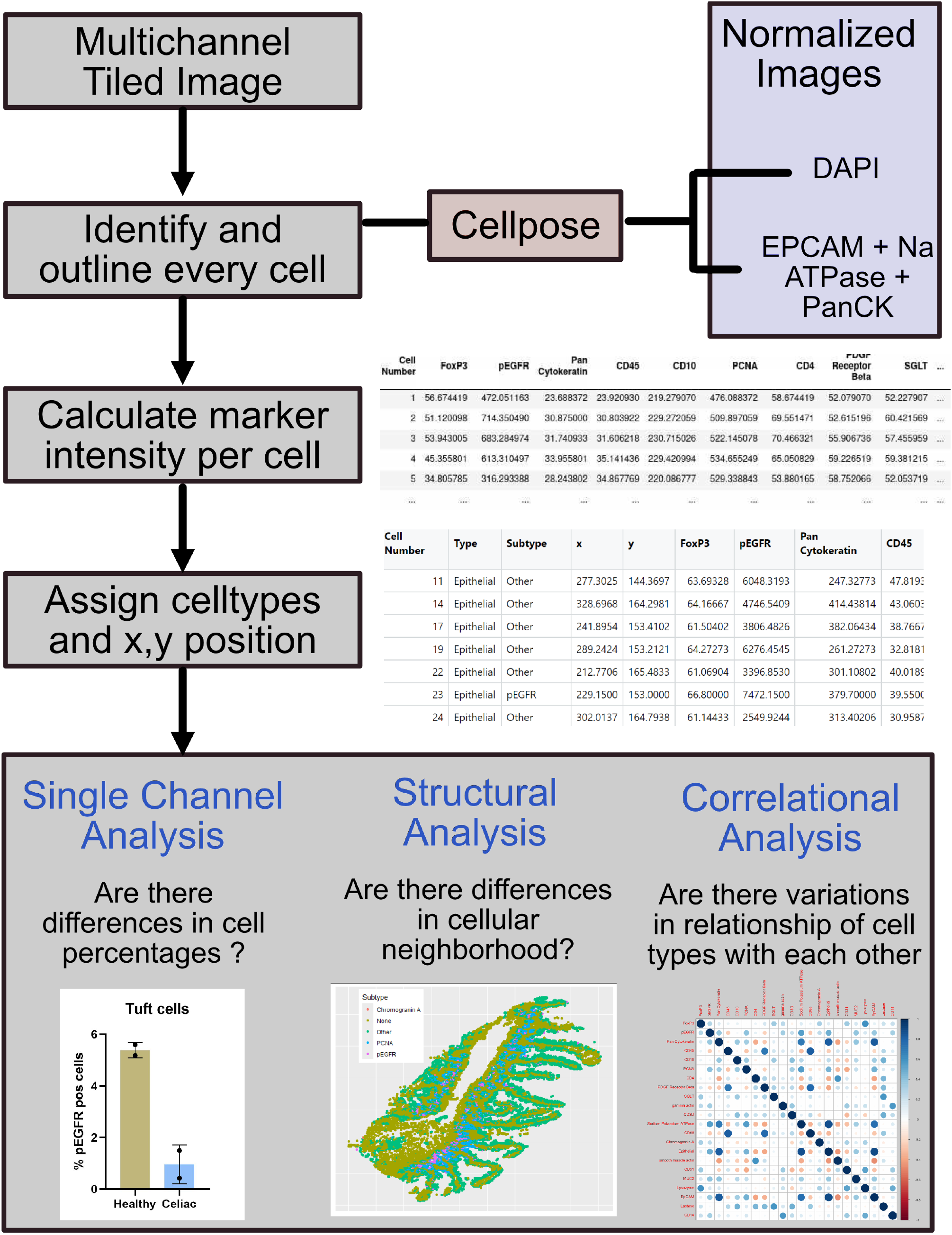
Schematic of analysis pipeline

As the first step in image analysis, a tissue mask was created. In a twofold downsampled image, all the individual images in the multichannel image were normalized and added to create a composite stain image. This was then thresholded by Otsu’s method to create a binary image which was then dilated. The largest contoured image with filled in holes was defined as the tissue mask. All subsequent image processing was conducted on images with the applied tissue mask.

Four channels (DAPI, Sodium Potassium ATPase, EpCAM, and Pan Cytokeratin) were normalized using Min Max Scaler in the sklearn python package. Sodium Potassium ATPase, EpCAM, and Pan Cytokeratin were then averaged together to create the final image for segmentation. CellPose (Ver 3.0.7) was then used for segmentation with the normalized cell boundary image and the normalized DAPI image being the “chan to segment” and “chan2” inputs respectively. The model used was “cyto3” with the default settings except for the “flow threshold” which was changed to 0.0.

**Figure S13:**
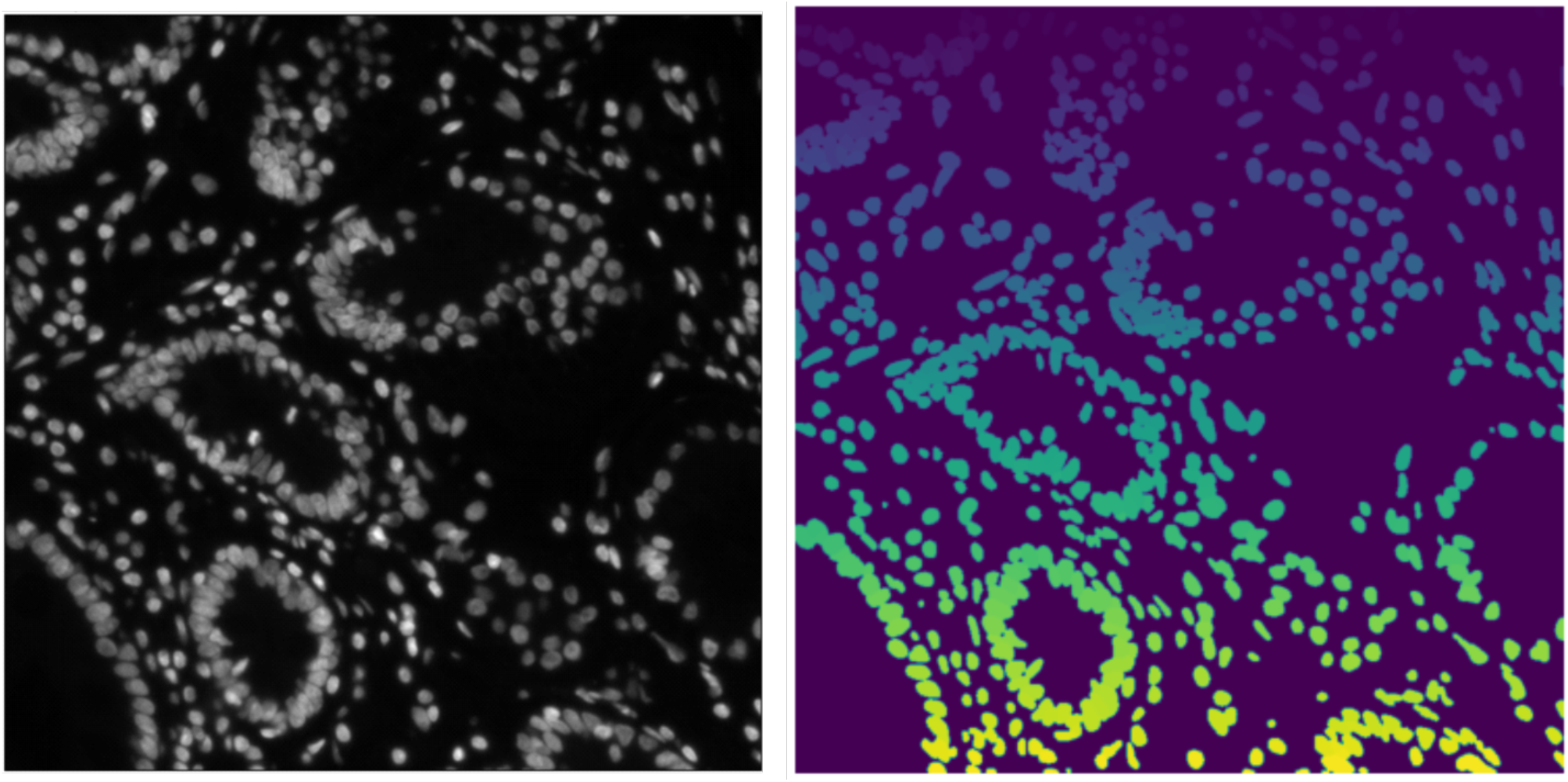
Cell nuclei segmentation

We observed that in a random field of view shown here, this approach had an accuracy of 86% with a precision of 94% when compared to a manual analysis.

The mask along with the tiled images are then used for further analysis using a custom python code. Data structure was created to determine the mean intensity of each protein, along with the centroid coordinates for each cell in the image. The celltypes were then assigned using a set of criteria for each marker.

### Supplementary 13: Example cost of a SPECTRE-plex run

| Antibody | Cost of Vial | Size of Vial (mL) | Size of Vial (uL) | Amount per run (uL) | Cost per run (\$) |
| --- | --- | --- | --- | --- | --- |
| Chromogranin A | 529 | 0.1 | 100 | 1 | 5.29 |
| sodium potassium ATPase | 610 | 0.1 | 100 | 2 | 12.2 |
| pEGFR | 565 | 0.1 | 100 | 5 | 28.25 |
| SGLT | 519 | 0.1 | 100 | 10 | 51.9 |
| Alexa Fluor™ 555 Antibody Labeling Kit | 136.8 | 0.1 | 100 | 10 | 13.68 |
| Pan CytoKeratin | 455 | 0.5 | 500 | 5 | 4.55 |
| FoxP3 | 165 | 0.125 | 125 | 10 | 13.2 |
| smooth muscle actin (SMA) | 590 | 0.1 | 100 | 2 | 11.8 |
| CD45 | 435 | 0.1 | 100 | 10 | 43.5 |
| CD14 | 625 | 0.1 | 100 | 10 | 62.5 |
| CD4 | 249 | 0.5 | 500 | 10 | 4.98 |
| CD68 | 435 | 0.1 | 100 | 10 | 43.5 |
| MUC2 | 610 | 0.1 | 100 | 0.5 | 3.05 |
| Lactase | 449 | 0.1 | 100 | 2 | 8.98 |
| CD3D | 600 | 0.1 | 100 | 5 | 30 |
| gamma actin | 350 | 1 | 1000 | 5 | 1.75 |
| CD10 | 350 | 0.1 | 100 | 10 | 35 |
| PDGF Receptor Beta | 217.5 | 0.05 | 50 | 5 | 21.75 |
| CD31 | 650 | 0.1 | 100 | 10 | 65 |
| PCNA | 350 | 1 | 1000 | 25 | 8.75 |
| EpCAM | 635 | 0.1 | 100 | 2 | 12.7 |
| Lysozyme | 499 | 0.1 | 100 | 5 | 24.95 |
| DAKO Diluent | 320 | 125 | 125000 | 2800 | 7.168 |
| Hoescht | 135 | 10 | 10000 | 0.2 | 0.0027 |
| PBS |  |  |  |  | 15.17 |
| **Total Cost** |  |  |  |  | **529.62** |

### Supplementary 14: Human tissue sections

De-identified formalin-fixed paraffin embedded duodenal tissue sections for multiplex staining were obtained under Boston Children’s Hospital IRB protocol #P00046566. Celiac disease (active disease, serologically positive) and age-matched healthy control sections were obtained from <12yo subjects.

---

**Figure S14:**
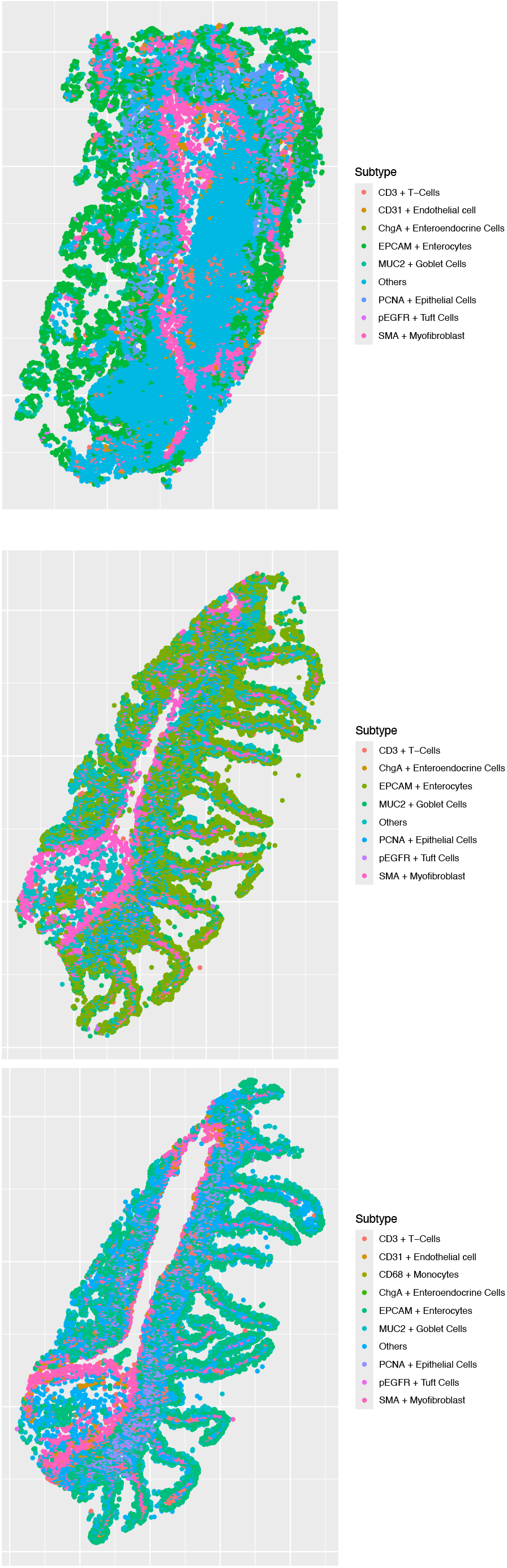
Spatial cell annotations for healthy tissues

**Figure S15:**
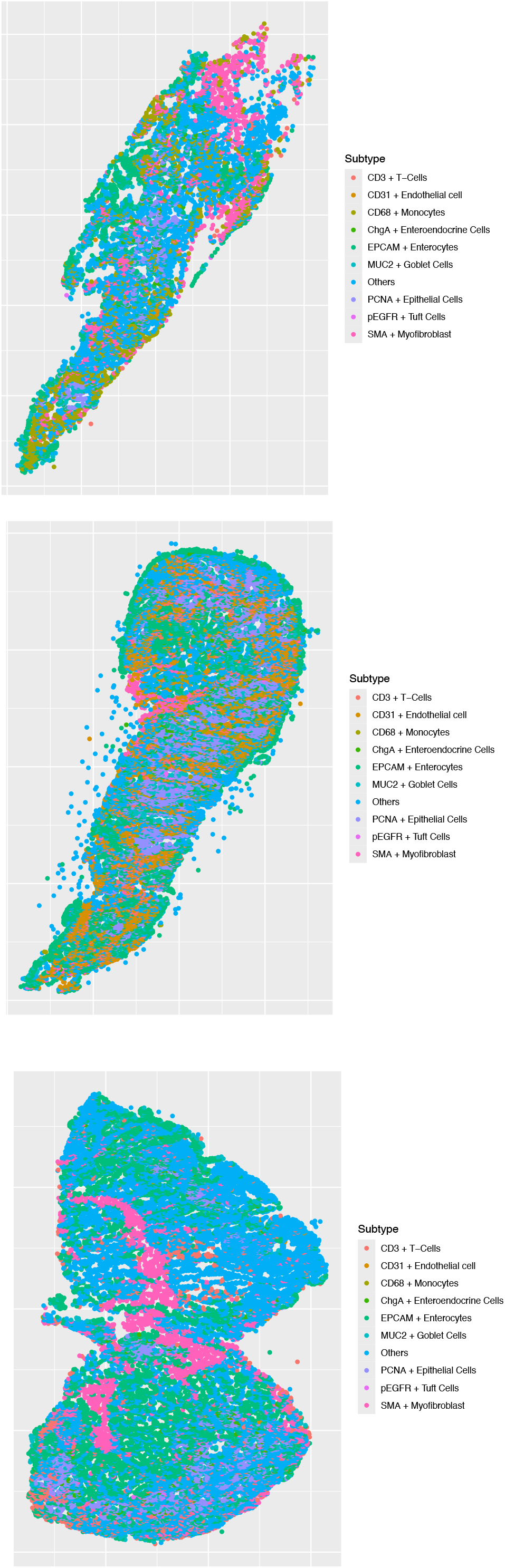
Spatial cell annotations for celiac disease tissues.

**Figure S16:**
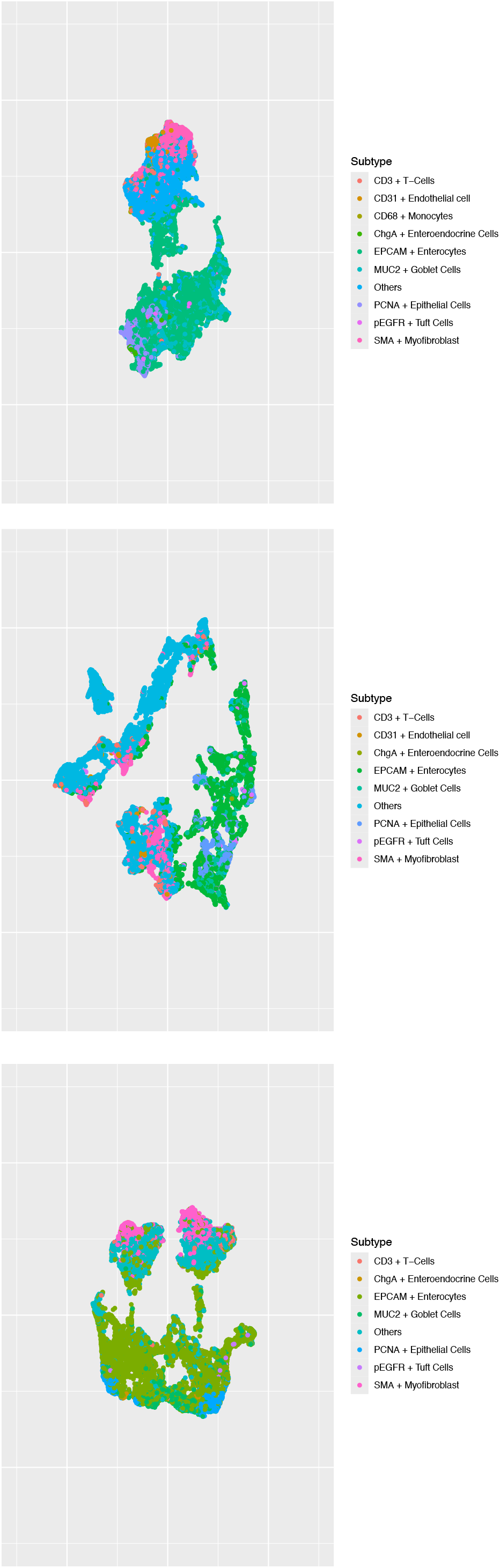
Uniform Manifold Approximation and Projection (UMAP) for spatial clustering of cell-types in healthy tissues

**Figure S17:**
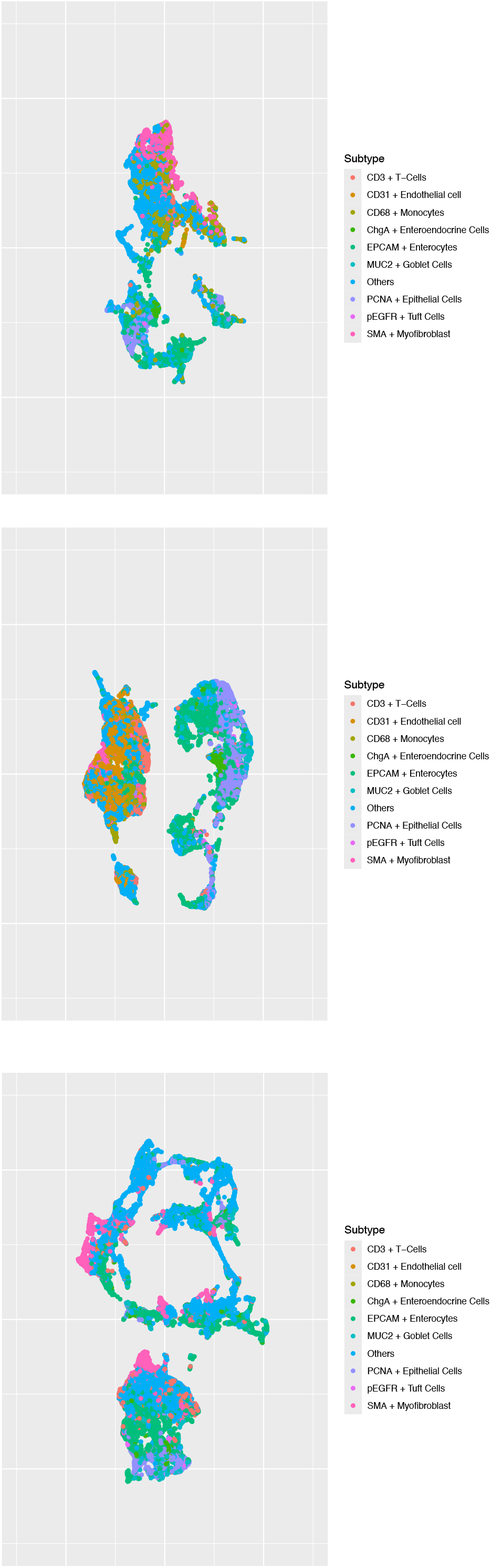
Uniform Manifold Approximation and Projection (UMAP) for spatial clustering of cell-types in celiac disease tissues

